# Met-HER3 crosstalk supports proliferation via MPZL3 in *MET*-amplified cancer cells

**DOI:** 10.1101/2021.07.27.454013

**Authors:** Yaakov E. Stern, Stephanie Duhamel, Abdulhameed Al-Ghabkari, Anie Monast, Benoit Fiset, Farzaneh Aboualizadeh, Zhong Yao, Igor Stagljar, Logan A. Walsh, Morag Park

## Abstract

Receptor tyrosine kinases (RTKs) are recognized as targets of precision medicine in human cancer upon their gene amplification or constitutive activation, resulting in increased downstream signal complexity including heterotypic crosstalk with other RTKs. The Met RTK exhibits such reciprocal crosstalk with several members of the human EGFR (HER) family of RTKs when amplified in cancer cells. We show that Met signaling converges on HER3 tyrosine phosphorylation across a panel of seven *MET*-amplified cancer cell lines and that HER3 is required for cancer cell expansion and oncogenic capacity in-vitro and in-vivo. Gene expression analysis of HER3-depleted cells identified MPZL3, encoding a single-pass transmembrane protein, as a HER3-dependent effector in multiple *MET*-amplified cancer cell lines. MPZL3 interacts with HER3 and MPZL3 loss phenocopies HER3 loss in *MET*-amplified cells, while MPZL3 overexpression rescues proliferation upon HER3 depletion. Together, these data support an oncogenic role for a HER3-MPZL3 axis in *MET*-amplified cancers.

## INTRODUCTION

Receptor tyrosine kinases (RTKs) have been among the earliest and most effective targets for precision medicine in human cancer (Yarden & Sliwkowski, 2001). RTK activation canonically proceeds through receptor homo-oligomerization and *trans*-autophosphorylation, but many cases of heterotypic signaling between different RTKs, frequently referred to as crosstalk, have been reported in the literature (Paul & Hristova, 2019; Ullrich & Schlessinger, 1990). Epidermal growth factor receptor (EGFR)-family RTKs in particular have become targets for clinical intervention in human cancer. The EGFR family contains four paralogous receptor tyrosine kinases that evolved from a single precursor EGFR homologue and exhibit extensive crosstalk with each other (Yarden & Sliwkowski, 2001). EGFR, human epidermal growth factor receptor 2 (HER2) and HER3 are frequently overexpressed in human cancers, and have been shown to induce canonical cancer-associated signals upon their activation by mutation, gene amplification or constitutive ligand presentation (Yamaoka, Kusumoto, Ando, Ohba, & Ohmori, 2018; Yarden & Sliwkowski, 2001). EGFR and HER2 have been successfully targeted both by ATP-competitive small molecule inhibitors as well as antibody-based therapies that bind to the extracellular domains of these RTKs. These interventions are part of the standard of care in lung, breast and colorectal cancer therapy (Khaddour et al., 2021; Oh & Bang, 2020).

Unlike EGFR and HER2, their paralogue HER3 does not possess intrinsic kinase activity due to substitutions at critical positions in the kinase domain (Knighton et al., 1993). In order to phosphorylate tyrosine residues in the HER3 cytoplasmic tail to recruit and activate intracellular signaling molecules, it must interact with another functional tyrosine kinase (Kovacs et al., 2015). HER3 exhibits this crosstalk preferentially with HER2 and EGFR, with which it can form heterodimers (Sliwkowski et al., 1994; Yarden & Sliwkowski, 2001). Ligand-induced heterodimerization with EGFR or HER2 promotes HER3 tyrosine phosphorylation at 6 positions (1054, 1197, 1222, 1260, 1276, and 1289) in the C-terminal tail capable of recruiting of the SH2-domain-containing p85 activator of phosphatidyl-inositol-3 kinase (PI3K) (Hellyer, Kim, & Koland, 2001; Schulze, Deng, & Mann, 2005) and promoting activation of the downstream Akt signaling pathway (Hellyer et al., 2001). Such activation of HER3 signaling by EGFR and HER2 is observed in lung and breast cancers, and combinatorial targeting of HER3 along with EGFR and HER2 via antibodies has been validated as a means to increase the efficacy of therapeutic intervention and delay or prevent the onset of acquired resistance (Gaborit et al., 2015; Junttila et al., 2009; Kodack et al., 2017).

The Met RTK, which has been assessed as a therapeutic target in multiple human solid tumours, has also been shown to engage in crosstalk with EGFR-family RTKs (reviewed in De Silva et al., 2017; Gherardi, Birchmeier, Birchmeier, & Vande Woude, 2012). *MET*-amplified cancer cell lines are exquisitely dependent on Met signaling for proliferation and survival through the activation of downstream mitogen-activated protein kinase (MAPK), PI3K-Akt and STAT3 signaling pathways (Lai et al., 2014; Smolen et al., 2006). Crosstalk between Met and EGFR family RTKs is observed in many cancer cell lines derived from *MET*-amplified lung, gastric, esophageal and other cancers (recently reviewed in Paul & Hristova, 2019). This has led to efforts to clinically target crosstalk between Met and EGFR itself, but despite combinatorial inhibition of Met and EGFR showing improved efficacy in cell culture and xenograft mouse models of non-small-cell lung cancer with *MET* amplification and *EGFR* mutations (Moores et al., 2016; Spigel et al., 2013; Xu et al., 2012; Y. W. Zhang et al., 2013), this benefit was not supported in clinical trials (Spigel et al., 2013). Nonetheless, *MET* amplification is recognized as a mechanism of resistance following small-molecule inhibitor targeting of EGFR mutant lung cancer, and conversely, EGFR and HER3 amplification or mutation have been observed as resistance mechanisms to Met inhibition in human cancers (Engelman et al., 2007; Gainor et al., 2016; Recondo et al., 2020). Conversely, ligand-mediated activation of EGFR-family RTKs, including HER3, restores cell proliferation in *MET-*amplified cells treated with a small-molecule Met inhibitor (Bachleitner-Hofmann et al., 2008). Thus, it remains critical to understand RTK crosstalk and its contribution to tumorigenesis as well as its role in the emergence of acquired resistance to RTK inhibitors in many cancer patients. Further elucidation of the mechanisms underlying RTK crosstalk downstream of Met will be important to improve patient stratification for the use of Met inhibitors and to effectively exploit co-vulnerabilities in Met and EGFR family signaling.

Here, we investigate crosstalk between Met and the EGFR family of RTKs using a panel of *MET*-amplified cancer cell lines as a model of Met-dependent cancers. We report selection of a Met-HER3 signaling axis across a panel of *MET*-amplified cancer cell lines that is required for cell proliferation. We identify a novel interactor of HER3, MPZL3, that is recruited in a Met-dependent manner and is required for efficient proliferation and tumour growth downstream of HER3 in *MET*-amplified cell lines. We further suggest that the HER3-MPZL3 axis may contribute broadly to RTK crosstalk in human cancer.

## RESULTS

### *Met crosstalk with the EGFR family converges on HER3 in* MET*-amplified cells*

To understand the crosstalk signaling axis between Met and EGFR family RTKs, we assembled a panel of *MET*-amplified cancer cell lines, derived from esophageal (OE33), gastric (KatoII, Okajima, Snu5, MKN45) and lung (EBC1, H1993) adenocarcinomas. SkBr3 cells were included as a control, in which EGFR, HER2 and HER3 phosphorylation is driven by *ERBB2* amplification and dependent on HER2 kinase activity (Figures 1C and S1). High levels of constitutively tyrosine phosphorylated Met protein were observed in OE33, KatoII, Okajima, Snu5, MKN45, EBC1 and H1993 cells as previously reported (Grandal et al., 2017; Lai et al., 2014; Tanizaki, Okamoto, Sakai, & Nakagawa, 2011), while SkBr3 cells did not express Met at a detectable level (Figure 1A). We assessed protein levels and tyrosine phosphorylation of EGFR family members EGFR, HER2 and HER3 by Western blot. Levels of the three EGFR-family receptors varies across the seven *MET*-amplified cell lines, although all three receptors are expressed and constitutively phosphorylated, comparable to the *ERBB2-*amplified SkBr3 cell line (Figure 1A). To establish whether EGFR, HER2 or HER3 phosphorylation is dependent on Met activity in our panel, we treated the panel of cell lines for one hour with PHA-665752 at (0.5 μM), a small-molecule tyrosine kinase inhibitor highly selective for Met over EGFR and HER2 at sub-micromolar concentrations (Christensen et al., 2003; Davis et al., 2011). While EGFR and HER2 were basally tyrosine phosphorylated in all cell lines tested, tyrosine phosphorylation of EGFR was dependent on Met kinase activity in 3 out of 7 tested cell lines (OE33, EBC1 and H1993) and tyrosine phosphorylation of HER2 was dependent on Met activity in two lung cancer cell lines tested (EBC1 and H1993) (Figure 1B). Tyrosine phosphorylation of HER3, by contrast to EGFR and HER2, was dependent on Met activity in all 7 *MET*-amplified cell lines tested (Figure 1B). This supports a preference for HER3 among EGFR family RTKs for Met dependent tyrosine phosphorylation.

**Figure 1:**
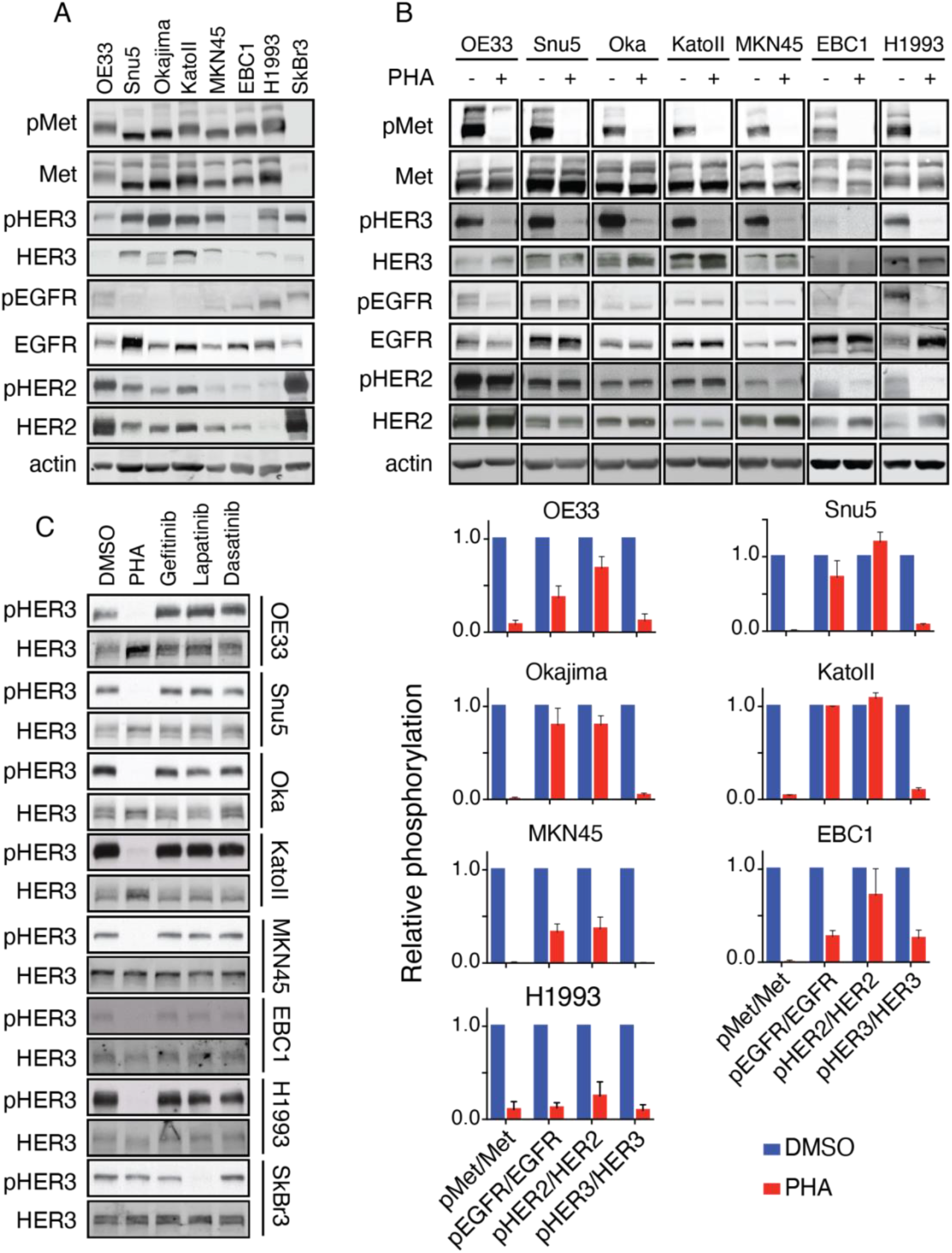
*MET*-amplified cancer cells display Met-EGFR family crosstalk with preference for HER3. (A) Western blot analysis of the phosphorylation of Met (Tyr^1234/45^), EGFR (Tyr^1173^), HER2 (Tyr^1221/22^) and HER3 (Tyr^1289^) in seven *MET*-amplified cancer cell lines and the *ERBB2*-amplified cell line SkBr3. (B) Western blot analysis of the phosphorylation of Met (Tyr^1234/45^), EGFR (Tyr^1173^), HER2 (Tyr^1221/22^) and HER3 (Tyr^1289^) in the indicated cell lines treated with PHA-665752 (PHA) (0.5 μM, 1 hour) or DMSO. Histogram shows quantification of phosphorylated RTK in treated vs. control conditions, normalized to total protein (lower panel, n=3). (C) Western blot analysis of the phosphorylation of HER3 (Tyr^1289^) in the indicated cell lines treated with PHA (0.5 μM); the EGFR inhibitor, gefitinib (1 μM); the HER2/EGFR inhibitor lapatinib (1 μM); or the Src and Abl family kinase inhibitor, dasatinib (0.1 μM), for 1 hour (n=3). Error bars indicate SEM.

As the HER3 kinase domain possesses impaired intrinsic tyrosine kinase activity, tyrosine phosphorylation of HER3 is normally attributed to its heterodimerization with EGFR or HER2 upon binding its ligand neuregulin, thus utilizing EGFR or HER2 kinase activity to induce HER3-dependent signaling (Yarden & Sliwkowski, 2001). While tyrosine phosphorylation of EGFR and HER2 remained mostly independent of Met activity in our panel of *MET*-amplified cell lines, it was possible that Met-dependent HER3 tyrosine phosphorylation also proceeded through an EGFR- or HER2-mediated mechanism. Additionally, Src family non-receptor tyrosine kinases are recruited to HER3 (Hause et al., 2012). Src overexpression has been shown to promote HER2-HER3 association (Ishizawar, Miyake, & Parsons, 2007), suggesting that Src could contribute to HER3 phosphorylation downstream of Met either directly or via HER2. Furthermore, Src can phosphorylate EGFR in the kinase activation loop, which potentiates EGFR kinase activity and signal output, indicating that Src could indirectly contribute to HER3 phosphorylation via an EGFR-dependent mechanism (Maa, Leu, McCarley, Schatzman, & Parsons, 1995).

To test whether HER3 phosphorylation proceeded through one of these mechanisms in *MET*-amplified cells, we treated the Met-dependent cell lines in our panel with PHA (0.5 μm); the EGFR small molecule kinase inhibitor, gefitinib (1 μm); the HER2 small molecule kinase inhibitor, lapatinib (1 μm); the Src-family broad-spectrum kinase inhibitor dasatinib (0.1 μm); or DMSO as control. Tyrosine phosphorylation of HER3 in all cell lines studied was decreased following inhibition of the Met kinase using PHA but not with any of the other inhibitors tested, indicating that Met kinase activity is primarily responsible for HER3 tyrosine phosphorylation in these *MET*-amplified cells (Figure 1C). Furthermore, the observation that HER2 phosphorylation was sensitive to lapatinib but not PHA in Snu5, Okajima, KatoII and OE33 cells indicates that the Met-HER3 signaling axis is separated from HER2 activity in these *MET*-amplified cancer cells (Figure S1). These data demonstrate that HER3 phosphorylation is primarily regulated by Met in a *MET*-amplified setting, and the consistent observation of this across seven independently derived cell lines further indicates that tyrosine phosphorylation of HER3 by Met is under strong selection in *MET*-amplified cancers.

### HER3 depletion impairs proliferation and reduces tumour growth in Met-dependent cancer cells

Recurrence of a Met-HER3 crosstalk axis across multiple *MET*-amplified cancer cell lines further suggests that HER3 may act as a ubiquitous signal transducer downstream from Met in this context. To test whether HER3 contributed to oncogenic transformation in *MET*-amplified cancer cells, we depleted HER3 using two independent shRNA hairpins in *MET-*amplified EBC1, H1993 and KatoII cells (Figure 2A). HER3 depletion decreased cell proliferation in all three *MET*-amplified cell lines tested when compared to controls (Figure 2B). HER3 knockdown also impaired the ability of KatoII cells to form colonies in soft agar, indicating that impaired cell expansion also affects oncogenic phenotypes canonically associated with Met-dependent transformation (Figure 2C) (Lai et al., 2014). In H1993 cells, which do not form colonies in soft agar under normal conditions (data not shown), HER3 knockdown impaired colony-forming capacity in 2D cell culture (Figure 2D). This stood in contrast to our observations in non-*MET-*amplified HeLa cells, in which HER3 knockdown did not exert any effect on clonogenic capacity in 2D cell culture (Figure S2A). This led us to test whether HER3 was required for *MET*-amplified tumour formation in-vivo.

**Figure 2:**
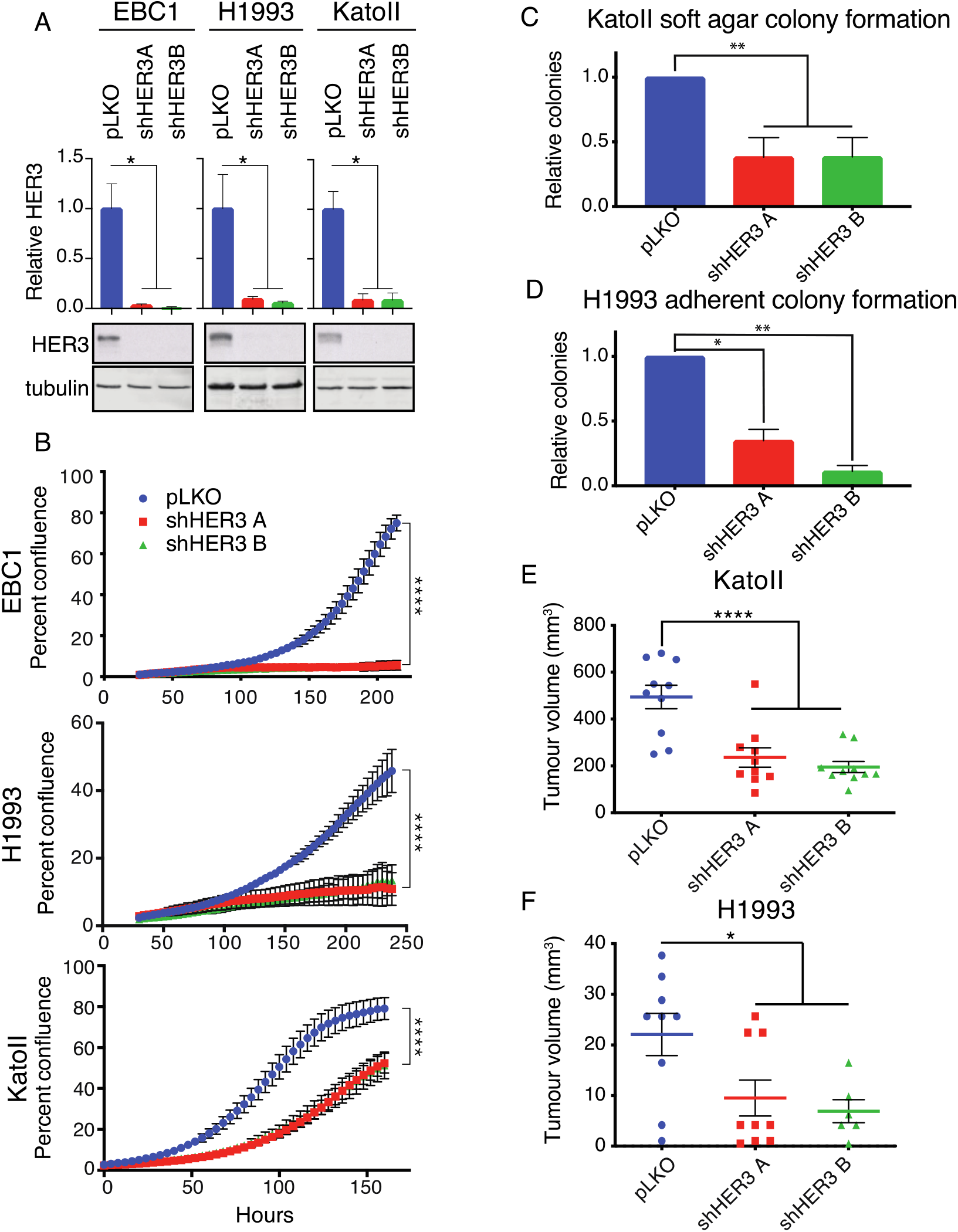
HER3 is required for expansion of *MET*-amplified cancer cells *in vitro* and *in vivo*. (A) Western blot analysis of total and phosphorylated levels of HER3 in the indicated cell lines stably transduced with control (pLKO) or shRNA targeting HER3. Quantification of HER3 relative to tubulin (n=3). (B) Live-cell imaging to measure cell confluence of the indicated cell lines stably transduced with control (pLKO) or shRNA targeting HER3. Representative replicates are shown (n=3); error bars indicate SD. (C) Quantification of colony-forming capacity of KatoII cells in soft agar (colonies >200 μm) (n=4). Error bars indicate SEM. (D) Quantification of colony-forming capacity in H1993 cells (n=3). Error bars indicate SEM. (E-F) Tumour growth of the indicated cell lines engraft in the flank of immunocompromised mice (NSG), n=10. One-way ANOVA with Dunnett’s test for multiple comparisons: *p≤ 0.05; **p ≤ 0.01; ***p≤ 0.001; ****p≤ 0.000.

To evaluate if HER3 depletion impacted tumorigenicity we subcutaneously injected immune-deficient NSG mice with H1993 and KatoII cells harbouring stable knockdown of HER3, or control cells infected with an empty shRNA expression vector (pLKO). Tumours formed rapidly from KatoII control cells, whereas tumours grown from HER3-depleted cells formed with delayed kinetics and were smaller than control tumours at all time points (Figure 2D and S2B). Tumours from HER3-depleted cells did not re-express HER3 protein, demonstrating that knockdown remained stable through the length of the experiment (Figure S2C). Subcutaneous tumours from H1993 control cells similarly grew more rapidly than those grown from HER3-depleted cells at early timepoints (43.1% [shHER3 A] and 20.9% [shHER3 B] of control, day 15) (Figure 2F). Our results demonstrate that knockdown of HER3 impaired outgrowth of *MET*-amplified xenograft tumours in-vivo.

### *Gene expression analysis of HER3-dependent transcripts across* MET *-amplified cell lines*

We predicted that depletion of HER3 would affect proliferation in *MET*-amplified cells by reducing the phosphorylation of signaling pathway proteins canonically activated downstream of Met and HER3. To test this, we analyzed protein extracts from HER3-depleted EBC1, H1993 and KatoII cells for phosphorylation of the activation loop residues (T202/Y204) on ERK1/2 for MAPK pathway activity and markers of mTORC2 activity (S473) on the Akt signaling transduction protein to test PI3K pathway activation. Phosphorylation of these molecules and the activation of their downstream transcriptional targets, along with tyrosine phosphorylation of the dimerization domain at position 705 in the transcription factor STAT3, have been shown to be critical for proliferation, colony formation and tumourigenesis in a panel of *MET*-amplified gastric cancer cell lines, including KatoII (Lai et al., 2014). However, we observed no significant change in these signaling pathways in HER3-depleted cells (Figure 3A). While Met-dependent HER3 signaling has been reported to suppress apoptosis, we did not observe a significant increase in apoptosis by annexin-V positivity under the same conditions (Figure S3). Despite this, HER3 depletion significantly reduced cell expansion in multiple cell culture conditions and across multiple Met-dependent cell lines. This supports a phenotype of HER3 depletion in *MET*-amplified cells that is not dependent on the inhibition of canonical HER3-associated signaling pathways, but on potential novel functions of HER3.

**Figure 3:**
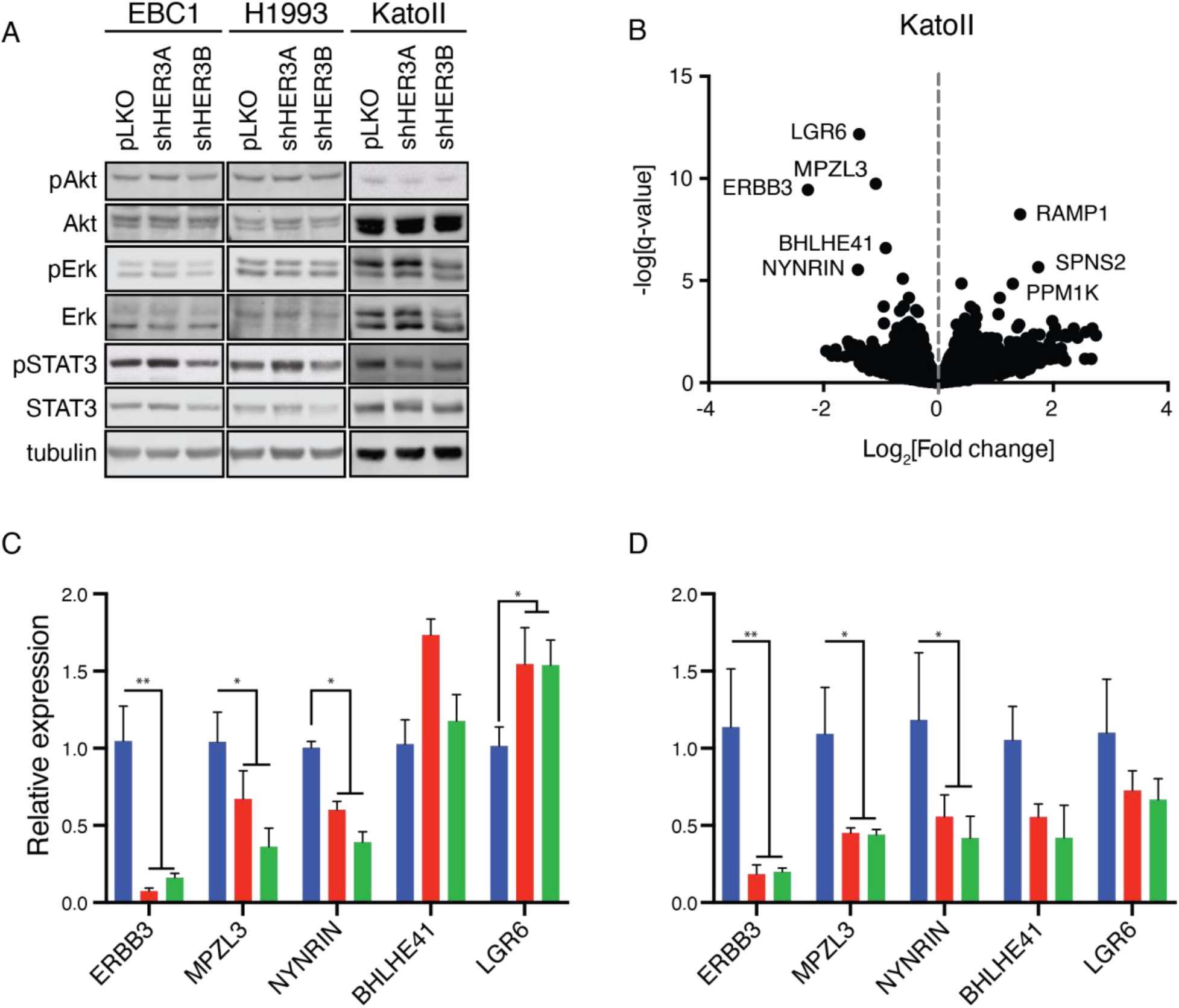
Gene expression analysis identifies MPZL3 as a HER3-regulated transcript in *MET*-amplified cells. (A) Western blot analysis of the phosphorylation of Akt (Ser^473^), ERK1/2 (Thr^202^/Tyr^204^) and STAT3 (Tyr^705^) in the indicated cell lines stably transduced with control (pLKO) or shRNA targeting HER3. (B) Volcano plot of RNA-sequencing data. Genes with significant increase or decrease in expression (log_2_FC ≥ |1|, FDR < 0.05) between control (pLKO) and HER3 depleted cells are labeled. (C-D) Measurement of the relative amounts of ERBB3, MPZL3, NYNRIN, BHLHE41 and LGR6 genes in the indicated cell lines by RT-qPCR. (C) Genes downregulated under steady state conditions upon stable HER3 depletion in EBC1 cells. (D) Genes downregulated under steady state conditions upon stable HER3 depletion in H1993 cells. Data are means (bars) of three individual replicates. Two-way ANOVA with Dunnett’s test for multiple comparisons; *p≤ 0.05; **p ≤ 0.01; ***p≤ 0.001; ****p≤ 0.0001. Error bars indicate SEM.

To identify HER3-regulated genes in *MET*-amplified cells we performed gene expression profiling by RNA sequencing of KatoII cells depleted of HER3 to identify significantly altered transcripts (FDR < 0.05, log_2_FC ≥ |1|). Genes with a decrease in abundance upon HER3 knockdown included ERBB3 as well as LGR6, MPZL3 and NYNRIN, while genes elevated in expression (log FC > 1, qval < 0.05) included PPM1K, RAMP1, and SPNS2 (Figure 3B). We expanded our analysis to include BHLHE41, a transcription factor involved in mammalian circadian regulation and a repressor of differentiation in skeletal muscle and the immune system (Ow, Tan, Jin, Bahirvani, & Taneja, 2014). BHLHE41 was significant in our analysis with a HER3-dependent fold change just below our cutoff (log_2_FC = 0.91, q-value = 8.42*10^−4^). To validate which transcripts were recurrently dependent on HER3 in a *MET*-amplified setting, we measured the expression of genes decreased in KatoII cells by quantitative RT-PCR in *MET*-amplified EBC1 and H1993 cells following HER3 knockdown.

Of the genes surveyed, ERBB3, MPZL3 and NYNRIN demonstrated reproducibly decreased mRNA levels in all HER3-depleted cells tested (EBC1, KatoII and H1993) (Figure 3C and D). LGR6 was unaffected and BHLHE41 was decreased in expression in H1993 and KatoII but not EBC1 cells. None of the genes elevated in expression upon HER3 knockdown in KatoII cells were differentially expressed in EBC1 or H1993 cells (Figure S4). NYNRIN is a gene with a putative RNA-binding function, but it has not been characterized biochemically to our knowledge (Mahamdallie et al., 2019). MPZL3 is a predicted adhesion receptor with a role in epidermal differentiation but no previously-characterized role downstream of RTKs (Cao et al., 2007; Wikramanayake et al., 2017). We proceeded to delineate the role of MPZL3 in Met-HER3 crosstalk in *MET*-amplified cells.

### *MPZL3 is required for proliferation downstream of HER3 in* MET *-amplified cells*

We tested the ability of MPZL3 to rescue HER3-dependent proliferation by overexpressing MZPL3 in EBC1 cells with or without knockdown of HER3. Empty-vector control and MPZL3-overexpressing EBC1 cells were then transfected with pooled siRNA targeting ERBB3 or control duplexes (Figure 4A). Cells were monitored for copy number for seven days, and HER3-depleted cells were compared to control-siRNA-treated cells for vector-control (pLKO) and MPZL3-overexpressing (MPZL3) cells, respectively (Figure 4B). Cells overexpressing MPZL3 were not sensitive to the diminished proliferation observed following to siRNA targeting of *ERBB3* (45.48% +/− 2.91%[SEM] vs 73.00+/− 3.11%[SEM] relative confluence; difference of means 27.53+/− 4.26%, 95% CI 19.07-35.98%; unpaired t test with Welch correction, Prism) (Figure 4B). This supports that MPZL3 overexpression can partially overcome the cell expansion defect following HER3 knockdown in *MET-* amplified cells.

**Figure 4:**
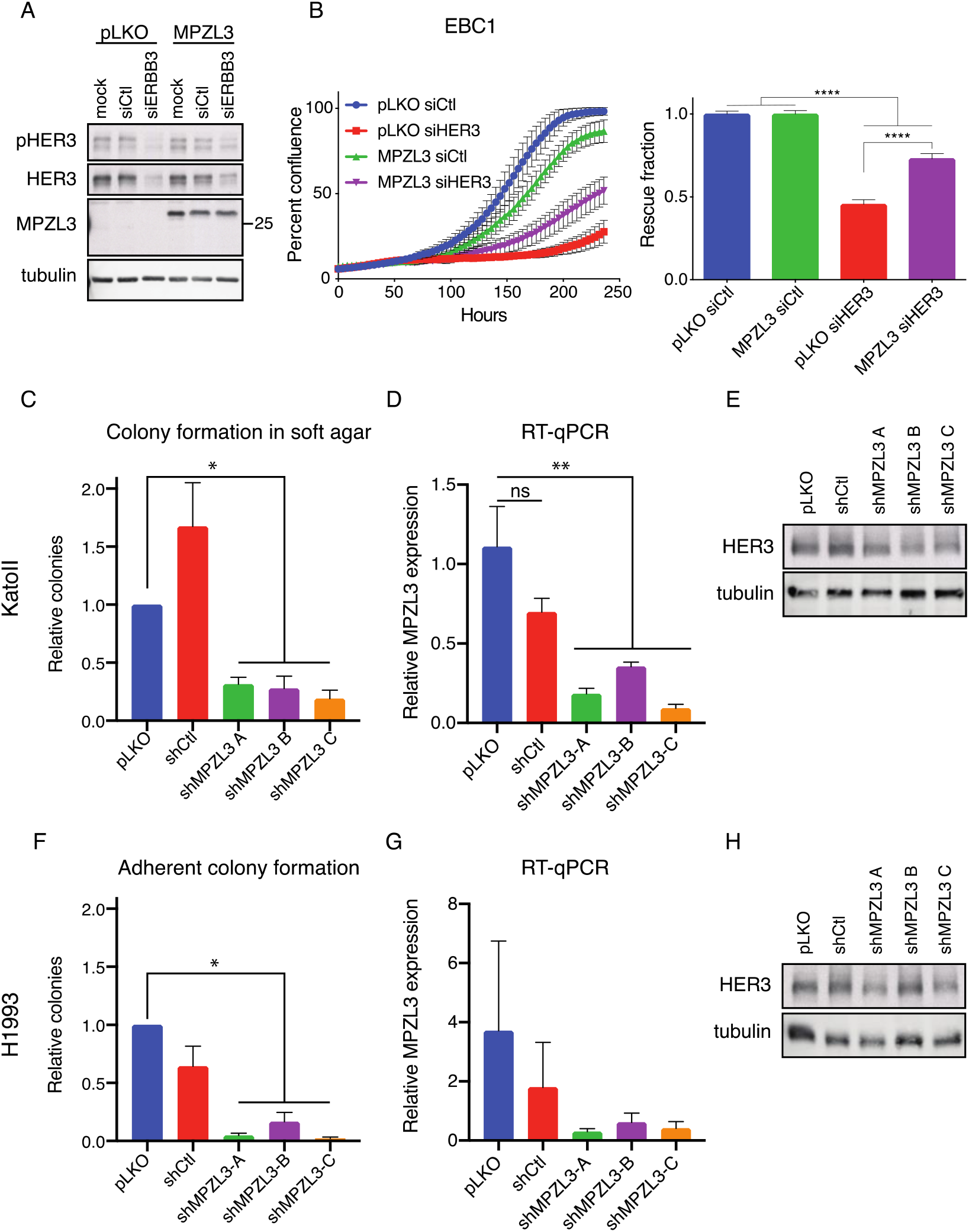
MPZL3 is required for *MET*-amplified cell expansion downstream of HER3. (A) Western blot analysis of total and phosphorylated levels of HER3 and MPZL3 in EBC1 cells overexpressing MPZL3, or an empty-vector control (pLKO) and transfected with control or HER3*-*targeting siRNAs. (B) Live-cell imaging to measure cell confluence of EBC1 cell lines stably transduced with control (pLKO) or MPZL3 and transfected with control or HER3*-*targeting siRNAs. Representative growth curves are shown; error bars show SD. Confluence relative to relevant siCtl-treated cells is quantified on the right panel (n=3); error bars show SEM. (C, F) Colony formation of KatoII cells in soft agar (C) or H1993 cells in adherent conditions (F) transduced with shRNA targeting MPZL3 or controls (pLKO and shCtl). Quantification of the colony number (n=4). (D, G) MPZL3 transcript levels in KatoII (D) and H1993 (G) cells transduced with shRNA targeting MPZL3 or controls (pLKO and shCtl). (E, H) Western blot analysis of total and phosphorylated levels of HER3 in KatoII (E) and H1993 (H) cells transduced with shRNA targeting MPZL3 or controls (pLKO and shCtl). (B): One-way ANOVA with the Sidak-Holm test for multiple comparisons; ****p≤ 0.0001. (C, D, F, G): One-way ANOVA with Dunnett’s test for multiple comparisons; *p≤ 0.05; **p ≤ 0.01.

To further test the contribution of MPZL3 to *MET*-amplified cells, we depleted MPZL3 using shRNA in cell lines in our panel. KatoII cells were impaired in their ability to form colonies in soft agar, as observed for HER3 knockdown (Figure 4C–E). We monitored MPZL3 depletion in H1993 *MET*-amplified cell lines by clonogenic assay under adherent conditions and again observed a dependence on MPZL3 similar to that seen for HER3 (Figures 4F–H). These data support a model whereby MPZL3 is critical for proliferation in *MET*-amplified cells, and that HER3-dependent stabilization of MPZL3 levels may underlie the requirement for HER3 for robust proliferation in *MET-*amplified cancers.

### HER3 and MPZL3 interact at the protein level

While our previous results demonstrate that HER3 depletion reduces the expression of MPZL3, we hypothesized that MPZL3 protein may interact with Met or HER3. To examine this, we transduced KatoII *MET*-amplified cells with lentiviral vectors encoding MPZL3-V5, GFP-V5 and an empty-vector control. We treated the cells for one hour with PHA to inhibit Met activation and then immunoprecipitated MPZL3 or GFP using an anti-V5 antibody (Figure 5A). HER3 co-immunoprecipitated with MPZL3-V5 in KatoII cells, but this interaction was reduced by Met inhibition (37.9% of DMSO control, +/− 11.2% [SEM.]). This supports the hypothesis that MPZL3 interacts with HER3, and that the HER3-MPZL3 interaction can be modulated by Met activity in *MET*-amplified cells.

**Figure 5:**
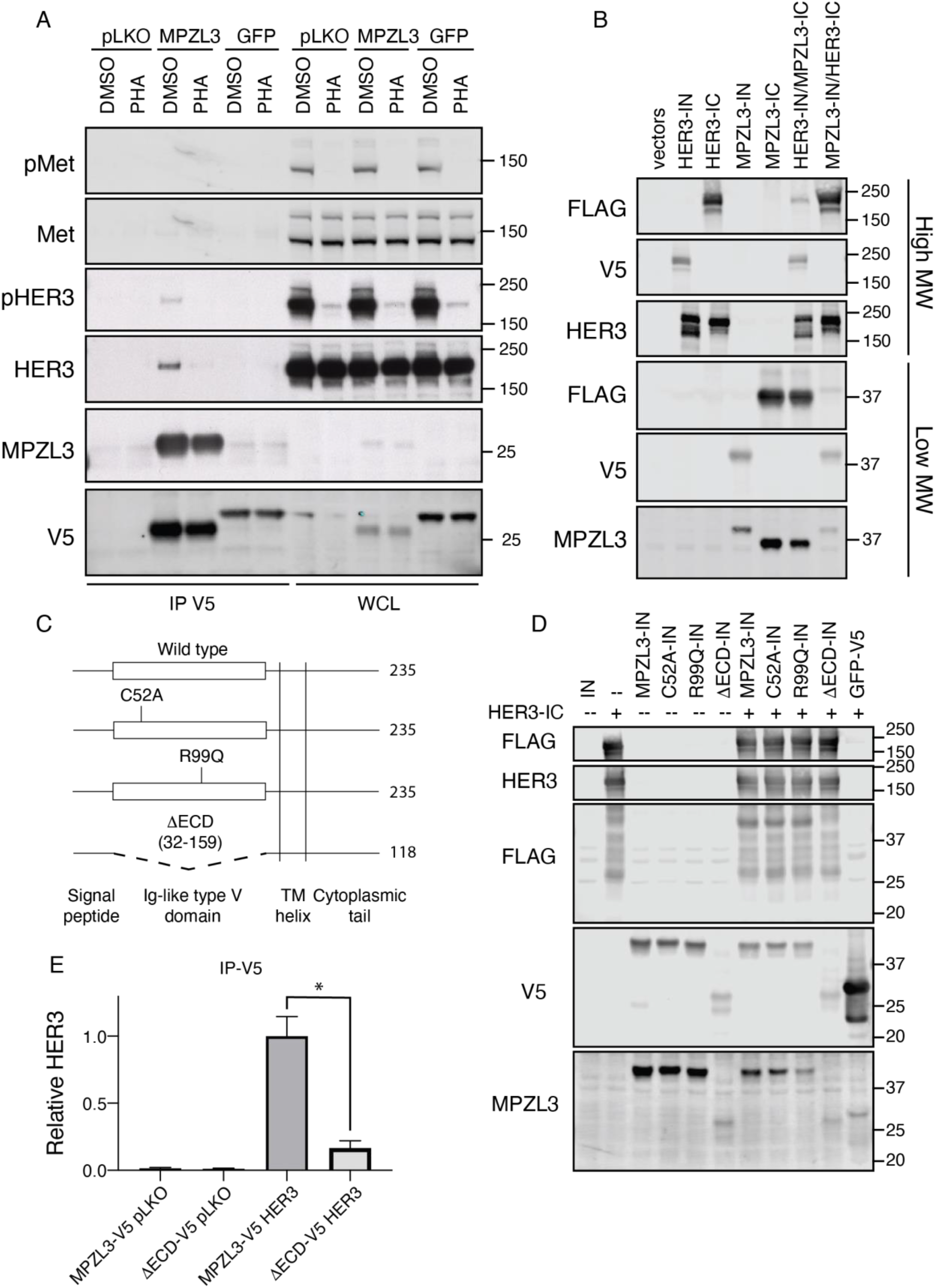
HER3 interacts with MPZL3. (A) Coimmunoprecipitation analysis of KatoII cells stably transduced with MPZL3- or GFP-V5 expression vectors, or control (pLKO). Cells were treated with PHA-665752 (0.5 μM, 1 hour) or with vehicle control (DMSO). (B) HER3 and MPZL3 were analyzed by split intein mediated protein ligation (SIMPL) assay to measure direct protein-protein interaction. A direct protein-protein interaction resulting in a reconstituted intein facilitates FLAG tag transfer from the IC-to the IN-linked fusion protein. (C) Mutants of MPZL3 predicted to impair the function of and a mutant deleting the extracellular domain (DECD) of MPZL3 were employed to uncouple the interaction between MPZL3 and HER3. (D) FLAG transfer by point mutants and the DECD mutant were compared to wild-type MPZL3 using the SIMPL assay. (E) Co-immunoprecipitation of MPZL3 and HER3 was quantified, comparing HER3 recovered by the full-length or DECD MPZL3 mutants. Data are means (bars) of three individual replicates. Student’s t-test; *p≤ 0.05; **p ≤ 0.01; ***p≤ 0.001; ****p≤ 0.0001. Error bars indicate SEM.

To test whether MPZL3 could interact directly with HER3, we employed the newly developed Split Intein-Mediated Protein Ligation (SIMPL) technique (Yao et al., 2020). SIMPL employs N- and C-terminal fragments of a self-splicing intein domain fused to bait and prey proteins. Upon interacting, the N- and C-terminal fragments reconstitute a full intein domain, splicing a FLAG peptide tag from the C-terminal intein domain (IC) to the N-intein-tagged (IN) prey protein (Yao et al., 2020). We cloned expression cassettes encoding HER3 and MPZL3 with IN- and IC-terminal intein fusions and co-expressed each pair in HEK293T cells. Co-expression of MPZL3 and HER3 induced fusion of the FLAG tag to the respective bait protein (Figure 5B). This indicated that MPZL3 and HER3 could interact directly in the absence of Met.

Next, we tested whether we could uncouple the HER3-MPZL3 interaction by mutating the single, extracellular Ig-like domain in MPZL3. We introduced point mutations in the extracellular domain predicted to impair the binding function of the Ig-like domain or deleted the entire Ig-like domain from MPZL3 (Figure 5C). Mutation of cysteine 52 to alanine (C52A) impairs the formation of a highly conserved disulfide bond characteristic of Ig-like domains and required for homophilic binding of the MPZL3 paralogue myelin protein zero (K. Zhang & Filbin, 1994). A spontaneous arginine-to-glutamine substitution in the extracellular domain of murine *MPZL3* (R99Q in human MPZL3) has been shown to induce progressive hair loss in mice, and this effect is phenocopied by homozygous deletion of *MPZL3*, indicating that this mutation impairs the major function of the gene under physiological conditions (Cao et al., 2007; Leiva et al., 2014; Wikramanayake et al., 2017). Using the SIMPL assay, we observed comparable FLAG transfer when wild-type, C52A or R99Q MPZL3 constructs were used as bait, but observed a loss of this effect upon deletion of the extracellular domain of MPZL3 (MPZL3-DECD) (Figure 5D). Deletion of the extracellular domain also strongly impaired co-immunoprecipitation of HER3 and MPZL3 upon their co-overexpression in HEK293T cells (16% of control) (Figure 5E). This supports a HER3-MPZL3 interaction dependent on the extracellular domain of MPZL3.

### *MPZL3 is overexpressed in* MET -, EGFR *- and* ERBB2 *-amplified cancer cell lines*

While we have observed a dependence on MPZL3 and its maintenance by HER3 in *MET*-amplified cells, it is unclear whether this is a specific requirement of Met-dependent transformation or whether MPZL3 expression is associated with RTK amplification in cancer. To understand if MPZL3 expression was generally altered in cancers with amplified RTKs, we interrogated the Cancer Cell Line Encyclopedia (CCLE) genomic copy-number and gene expression data to explore the relationship between RTK amplification and MPZL3 expression (Ghandi et al., 2019). First, we identified *MET*-amplified cell lines in the CCLE by analysis of relative copy number of the *MET* gene. Met amplification has been observed to correlate with response to Met-targeted therapy in patients with a *MET:CEP7* ratio of 5 or greater, indicating at least a 5-fold increase in *MET* copy number above ploidy (Camidge et al., 2014; Garber, 2014). CCLE data is reported from whole exome sequencing reads normalized to relative overall ploidy. Cell lines with *MET* copy relative to ploidy + 1 ≥ 2.3 would thus be predicted to have a *MET:CEP7* ratio of ≥ 5. We profiled all 58 RTK genes and annotated each cell line for amplification for every RTK gene. Cells with five or more copies of any RTK (copy relative to ploidy +1 ≥ 2.3) were considered amplified for that RTK. Of 1280 cell lines with data available, 98 (7.8%) had at least one RTK gene amplified. None of the cell lines with amplification of an RTK also showed amplification of the *MPZL3* gene at this threshold, indicating that the relationship of MPZL3 with an amplified receptor may be primarily under transcriptional regulation. We identified eleven cell lines as *MET*-amplified by our criteria (Table S1). These cell lines overexpressed MPZL3 at the transcript level relative to cell lines with no RTK amplified (median MPZL3 TPM 2.3 vs. 1.7, p=0.0306 [two-tailed Mann-Whitney test]) (Figure 6A). Next, we analyzed all amplified RTKs to determine whether transcriptional elevation of MPZL3 might be associated with other RTKs. Comparing across all RTKs amplified in multiple CCLE cell lines, *EGFR* and *ERBB2* amplification significantly predicted overexpression of MPZL3 relative to cells with no amplified RTK (mean MPZL3 TPM 2.83 ± 0.29 [EGFR] and 3.07 ± 0.24 [ERBB2] vs. 1.81± [no RTK]) (Figure 6B). EGFR and HER2, the *ERBB2* gene product, are known to heterodimerize with HER3, and both EGFR- and HER2-dependent cancer cell lines have been shown to have impaired proliferation upon the inhibition or depletion of HER3 (Gaborit et al., 2015; Junttila et al., 2009). Thus, the CCLE copy number and gene expression data support elevation of MPZL3 mRNA levels downstream of multiple amplified RTKs in human cancer cell lines.

**Figure 6:**
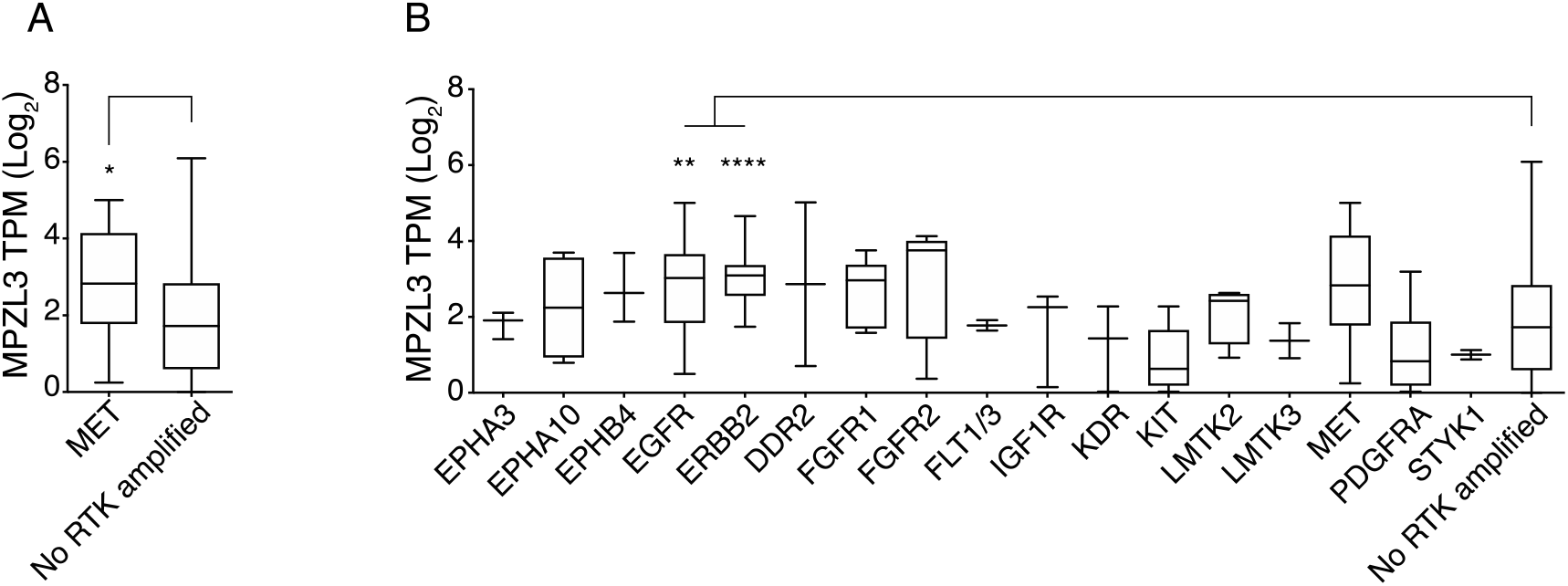
Amplification of genes encoding Met, EGFR and ERBB2 correlate with elevated levels of MPZL3 in the Cancer Cell Line Encyclopedia. (A) Met-amplified cells were identified in the CCLE by relative gene copy number (Log_2_ relative to ploidy +1 ≥ 2.3), and MPZL3 expression was observed to be higher than that in cells with no RTK amplified (median TPM: 2.8 [*MET*-amplified] vs. 1.7 [control], Mann-Whitney p=0.03). (B) Cell lines in CCLE were profiled for RTK amplification (Log_2_ relative to ploidy +1 ≥ 2.3), and MPZL3 expression was compared in cells with an amplified RTK to cells with no RTK amplification. ANOVA p<0.0001).

## DISCUSSION

Cancers harboring high-level amplification of the *MET* gene, with more than 5 copies of *MET* for each copy of chromosome 7, have been known to enrich for response to Met inhibitors in the clinic (Lennerz et al., 2011; Noonan et al., 2016; Ou et al., 2011; Tong et al., 2016). *MET* amplification and overexpression correlates with constitutive activation of the Met RTK and downstream signaling pathways in the absence of the Met ligand, HGF, in cancer-derived cell lines, indicating that the induction of constitutive Met signaling underlies this clinical observation (Lai et al., 2014; Smolen et al., 2006). By analyzing preference within the EGFR family for crosstalk with Met in *MET*-amplified cell lines, we have uncovered a critical role for HER3 in Met-dependent oncogenic signaling and have shown that this involves the expression and recruitment of a novel HER3-dependent transcript and interactor, MPZL3.

We found that while EGFR, HER2 and HER3 were co-expressed and usually tyrosine phosphorylated in our panel of *MET*-amplified cell lines, HER3 phosphorylation consistently showed dependence on Met activity, while EGFR and HER2 were frequently phosphorylated independently of Met. Supporting our biochemical observations that a Met-HER3 axis recurrently arises in *MET*-amplified cells, we found that HER3 expression was important for core oncogenic processes downstream of Met in three independent *MET*-amplified cell lines. Collectively, these results demonstrate a convergent selection for crosstalk between HER3 and the Met signaling pathway in Met-dependent cancers. Furthermore, our analysis of upstream kinases shows that while HER2 kinase activity is required for HER2 phosphorylation in most *MET*-amplified cell lines, this has no impact on HER3 phosphorylation, indicating co-option of HER3 signaling by Met in the presence of an intact HER2 signaling module. This suggest that a HER3-dependent signaling axis plays an important role in *MET*-amplified cells and is under strong selection for Met-dependent activation in this context.

HER3 has been shown to play a critical role in EGFR- and HER2-dependent cancers, including cancer cells harboring an EGFR mutation or amplification of the *ERBB2* gene (Engelman et al., 2007; Junttila et al., 2009; Yonesaka et al., 2015). Intriguingly, in EGFR-mutant HCC827 lung cancer cells subjected to prolonged gefitinib treatment in tissue culture, an acquired resistance mechanism involved *MET* gene amplification, Met-dependent HER3-tyrosine phosphorylation and HER3-dependent activation of PI3K downstream of Met (Engelman et al., 2007). *MET* amplification has been observed lung cancer patients treated with the EGFR inhibitors gefitinib and erlotinib, and treatment of gefitinib-resistant tumours with the Met inhibitor crizotinib is effective in the clinic for these patients (Gainor et al., 2016). While this has been suggested to depend on activation of the PI3K signaling pathways including the Akt pathway, in our panel of *MET* amplified cells, PI3K activity-associated phosphorylation of Akt remained intact following inhibition of Met kinase with small molecule inhibitors. We similarly did not observe a significant increase in the apoptotic marker annexin-V in EBC1, H1993 and KatoII *MET*-amplified cells upon depletion of HER3, or changes in known Akt-dependent gene expression (data not shown), supporting the idea that Met-dependent activation of pro-survival signaling downstream of the Akt pathway remained intact. This, in turn, indicated that the contribution of HER3 to Met-dependent signaling proceeded through a novel HER3-dependent mechanism, and did not impact the survival pathways canonically associated with oncogenic HER3 signaling.

The highest-confidence and most differentially expressed candidate genes downstream of HER3 were genes whose expression decreased upon HER3 knockdown, including *MPZL3*, a gene encoding a transmembrane protein with an extracellular Ig-like domain and an unstructured cytoplasmic tail involved in epidermal differentiation (Racz et al., 2009; Wikramanayake et al., 2017). Our observations that suppression of MPZL3 phenocopies HER3 knockdown and that MPZL3 overexpression can partially rescue proliferation upon HER3 knockdown support a role for MPZL3 downstream of HER3 in *MET*-amplified cell lines. This would explain the recurrence of Met-HER3 crosstalk in *MET*-amplified cells, as a proliferative advantage would lead to convergent selection for this signaling axis.

MPZL3 was initially identified as a regulator of skin morphogenesis via a spontaneous point mutation, leading to an arginine-to-glutamine (R99Q in human MPZL3) amino acid substitution in the extracellular Ig-like domain and a progressive hair loss phenotype in laboratory mice (Cao et al., 2007). This is due to impaired differentiation of sebaceous gland precursors in MPZL3-mutant or -knockout mice (Cao et al., 2007; Leiva et al., 2014; Wikramanayake et al., 2017). Our co-immunoprecipitation and SIMPL assay experiments demonstrate that the extracellular Ig-like domain is required for the formation of a HER3-MPZL3 complex. The C52A and R99Q mutations, however, do not impair the association of MPZL3 with HER3, implying that different functions of the Ig-like domain are required for MPZL3-dependent epidermal differentiation and its interaction with HER3.

Our observations suggest that *MET* amplification co-opts HER3 and its binding partner MPZL3 to support oncogenic cell proliferation via a previously-unreported mechanism. This was supported by our observation that *MET* gene amplification predicted higher *MPZL3* expression than cell lines without any RTK amplification in the CCLE. While *MPZL3* has been reported to act as a tumour suppressor in cutaneous squamous cell carcinoma through its activity promoting differentiation (Bhaduri et al., 2015), our proliferation and colony-forming assays demonstrate an oncogenic role for *MPZL3* in lung and gastric carcinoma cells. Convergence on a shared HER3-MPZL3 dependency could explain the observation that Met and the EGFR family receptors are important bypass pathways for each other in a number of models of resistance to EGFR or HER2 inhibition, as reactivation of any of these receptors could stabilize MPZL3 levels in a similar manner (Bachleitner-Hofmann et al., 2008; Cappuzzo et al., 2009; Engelman et al., 2007; Gainor et al., 2016; Recondo et al., 2020). As MPZL3 and HER3 interact directly, it may be possible to identify cancers dependent on this interaction and the upstream activity of HER3-activating kinases such as Met, EGFR and HER2 via this mechanism. Intriguingly, HER3 phosphorylation has been repeatedly observed as a bypass mechanism of resistance to BRAF and MEK1/2 inhibitors in melanoma, indicating that there may be additional mechanisms in which HER3 plays a critical and central role in acquired resistance (Abel et al., 2013; Capparelli et al., 2018; Cheng et al., 2015; Han et al., 2018). Further analysis of the emerging biology of HER3 and MPZL3 and their activities in human cancer will be critical to fully exploiting their potential in cancer therapy.

## MATERIALS AND METHODS

### Antibodies and reagents

Antibody 148 was raised in rabbit against a C-terminal peptide of human Met (Rodrigues, Naujokas, & Park, 1991). Antibodies against phosphorylated Met at Tyr^1234/1235^, HER2 at Tyr^1221/1222^, HER3 at Tyr^1289^, Akt at Ser^473^, Erk1/2 at Thr^202^/Tyr^204^, STAT3 at Tyr^705^, and generic tyrosine peptide (pY100), as well as total EGFR, HER2, HER3, pan-Akt, Erk1/2, and STAT3 were purchased from Cell Signaling Technologies. Antibodies against phosphorylated EGFR at Tyr^1173^ were purchased from Santa Cruz Biotechnology. Antibodies against M2-FLAG peptide, actin and tubulin were purchased from Sigma. Antibodies against V5 peptide were purchased from Abcam, while antibodies against MPZL3 were purchased from ProteinTech. HRP-conjugated secondary antibody was purchased from Thermo Fisher, while IRDye infrared secondary antibodies were purchased from Mandel Scientific.

PHA-665752 (Met inhibitor) was used at 0.5 μM and was a gift from Pfizer. Gefitinib and lapatinib, used at 1 μM, and dasatinib, used at 100 nM, were gifted by Dr. W. Muller. Dimethyl sulfoxide (DMSO) was purchased from Sigma and used at a concentration of 1:1000 as a vehicle control.

### Cell cultures and RNAi

OE33 cells were described previously (Grandal et al., 2017) and cultured in RPMI supplemented with 10% FBS and 2 mM L-glutamine (Thermo Fisher). Snu5, KatoII, Okajima and MKN45 cells were cultured as described previously (Abella, Parachoniak, Sangwan, & Park, 2010; Lai et al., 2014; Lai, Durrant, Zuo, Ratcliffe, & Park, 2012). EBC1 were obtained through RIKEN BRC Cell Bank and H1993 cells were obtained from ATCC; these were cultured in MEM alpha or RPMI media, respectively, supplemented with 10% FBS (Thermo Fisher). SkBr3 cells were gifted by Dr. William Muller and were cultured in McCoy’s 5A media supplemented with 10% FBS (Themo Fisher). HEK293T cells were cultured in DMEM supplemented with 10% FBS (Thermo Fisher).

EBC1 cells were transfected in suspension using pooled siRNA at 50 nM using the HiPerfect protocol (Qiagen) and immediately seeded for proliferation and lysis, which was conducted after 72 hours. Allstars control siRNA was purchased from Qiagen. siGENOME SMARTpool pooled siRNA duplexes targeting *ERBB3* were purchased from Dharmacon (sequences in Supplementary Table 1). Viral vectors for shRNA expression as well as empty-cassette plasmids used as vector controls were obtained from the Mission TRC library (Sigma) via the McGill Platform for Cellular Perturbation (clone information and sequences in Supplementary Table 2). Gateway entry vector for MPZL3 (clone information and sequence in Supplementary Table 3), Gateway destination vectors pLX303, pLX304, and pLEX307 for mammalian cDNA overexpression, and pLX317 vector for GFP-V5 overexpression were obtained through the Mission TRC3 Orfeome collection via the McGill Cellular Perturbation Services. Split intein mediated protein ligation (SIMPL) Gateway destination vectors were generously gifted by the Stagljar lab. Gateway entry vector for ERBB3 was obtained through Addgene (pDONR223-ERBB3, #23874) (Johannesen *et al.*, 2010). Vector recombination was performed using the Gateway LR Clonase II enzyme kit (Thermo Fisher).

### Lentiviral transduction

Transfections of HEK293T cells for viral production were performed by the calcium phosphate method. Lentiviral cultures for shRNA or transgene overexpression were collected as described at https://portals.broadinstitute.org/gpp/public/resources/protocols. Lentiviral transduction was followed by selection in puromycin (1-2 μg/ml) or blasticidin (2-10 μg/ml) for 3-14 days, and control cells were verified for negative selection prior to the start of each experiment.

### Proliferation and soft agar experiments

Proliferation assays were performed in 96-well dishes with 4×10^3^ EBC1, H1993 or KatoII cells seeded per well with 18 replicates per condition. Proliferation was measured as a function of cell confluence by IncuCyte live-cell microscopy (Essen Biosciences). Soft agar assays were performed in 6-well dishes coated with a layer of bottom agar (6 mg/ml), with 1×10^4^ KatoII cells seeded in top agar (3.2 mg/ml). Cells were plated in duplicate per condition and fed every 4-5 days, with colonies counted after 23 days in culture. For both proliferation and soft agar assays, representative examples are shown of three independent replicates. Quantification was performed from all three replicates where shown.

### Subcutaneous xenograft experiments

For xenograft experiments, 5×10^5^ (KatoII) or 1×10^6^ (H1993) cells were injected bilaterally into the flanks of immune-deficient NOD.Cg-Prkdc^scid^Il2rg^tm1Wjl/SzJ^ mice (Jackson). Five mice were injected per condition. Tumours were detected by palpation and measured every 3 days. Animals injected with KatoII cells were sacrificed for ethical considerations at 13 days post-injection.

### Flow cytometry

EBC1, H1993 or KatoII cells selected for virally-transduced shRNA vectors (puromycin) were seeded in equal numbers and grown for 3 days before washing in PBS, followed analysis of cell surface annexin-V and cell permeability using the Annexin-V-FLUOS staining kit (Sigma). Flow cytometry was performed on live cells using a BD FACSCalibur cytometer at the McGill Life Sciences Complex Flow Cytometry Facility.

### Immunoprecipitation and Western blotting

Cells were harvested for protein lysis in a Triton-glycerol-HEPES (TGH) buffer described previously (Lai et al., 2012). Lysates were frozen, thawed and centrifuged at full speed in an Eppendorf microcentrifuge for 15 minutes and protein content was assessed by Bradford assay (Biorad). Cell lysate equivalent to 1 mg protein collected 36 hours post-transfection was diluted to 500 μl in TGH buffer for each immunoprecipitation reaction and precleared with 10 μl sepharose-protein G beads (GE Healthcare). Immunoprecipitation was performed overnight by incubation with anti-V5 antibody, followed by capture using sepharose-protein G beads. Beads were washed three times in lysis buffer prior to analysis.

Centrifuged whole cell lysate or proteins captured on sepharose beads were boiled for 5 minutes in Laemmli buffer. Samples were resolved using 8% or 12% SDS-PAGE (BioRad) or 4-12% gradient NuPAGE Protein Gel (Thermo Fisher). Proteins were transferred to Newblot PVDF membranes (Mandel Scientific), blocked in 3% bovine serum albumin in TBST or Odyssey Blocking Buffer (Licor) and incubated with primary antibodies diluted in Near Infra-Red Blocking Buffer (Rockland) overnight at 4° C. Membranes were washed in TBST and incubated with secondary antibodies diluted in a 1:1 TBST:Rockland blocking buffer for one hour, washed in TBST again, and scanned using a Odyssey scanning machine (Licor) or visualized using Western Lightning Plus ECL (Perkin Elmer).

### RNA sequencing

KatoII cells transduced with shHER3A, shHER3B or the pLKO empty vector were selected in puromycin and plated for experiments. Total RNA was extracted from 3 independent infections used to prepare replicates for soft agar and xenograft injection assay using the RNeasy Plus Mini Kit (Qiagen). High RNA quality in all samples was verified using the Bioanalyzer RNA Nano 600 assay (Agilent). cDNA libraries were PCR-amplified and barcoded from 500 nanograms of RNA per sample, and pooled libraries were sequenced using the HiSeq4000 next-generation sequencing platform (Illumina) by the Centre d’Expertise et de Services at Génome Québec. The pooled libraries were sequenced with an average coverage of 3.3×10^7^ reads per sample.

Fastp (version 0.19.4) was used to collect QC metrics of the raw reads. RNA sequences were aligned and sorted by coordinates, to the NCBI human genome build 38 (GRCh38.p12) with version 94 of gene annotations, using the STAR aligner (STAR_2.6.1a_08-27) (Chen, Zhou, Chen, & Gu, 2018; Dobin et al., 2013). The removal of alignment duplicates was done with Sambamba (version 0.6.8) (Tarasov, Vilella, Cuppen, Nijman, & Prins, 2015). Quantification of genes was performed using featureCounts (v1.6.3) (Liao, Smyth, & Shi, 2014). DESeq2 (v 1.20.0) was used to normalize feature counts and to test find the differentially expressed genes (Love, Huber, & Anders, 2014). The HGNC symbols were extracted and added to the DESeq2 results data frame using biomaRt (version 2.36.1) using the “hsapiens_gene_ensembl” dataset and the “Ensembl Release 94 (October 2018)” mart (Durinck et al., 2005; Durinck, Spellman, Birney, & Huber, 2009).

### Quantitative RT-PCR

RNA was isolated from three independently infected and selected lines of EBC1 and H1993 cells using the RNeasy Plus Mini Kit (Qiagen). High RNA quality in all samples was verified using the Bioanalyzer RNA Nano 600 assay (Agilent). cDNA libraries were synthesized using the Transcriptor First Strand cDNA Synthesis Kit (Roche). qPCR reactions were performed using SYBR Green I Master on a LightCycler480 (Roche). Gene expression was normalized to the *GAPDH, HPRT* and *RPLP0* genes using the delta-delta-Ct method. Primers for qPCR were purchased from Integrated DNA Technologies (primer information listed in Supplementary Table 4).

### Statistical Analysis

Quantitative data are presented as the means ± SEM. Statistical significance was assessed using a two-tailed Student’s *t* test, and ordinary one-way ANOVA with Tukey’s correction for multiple comparisons, unless otherwise indicated, using Prism software. Significance is as follows: p > 0.05, not significant (ns); *p ≤ 0.05; **p ≤ 0.01; ***p ≤ 0.001; ****p ≤ 0.0001. Data distribution was assumed to be normal, but this was not formally tested. P values and the number of experiments used for quantification and statistical analysis are indicated in the corresponding figure legends.

## Data availability

GEO reference for RNA-Seq: *Deposit process ongoing.*

## ACKNOWLEDGEMENTS

We thank Dr. Phillippe Daoust and Dr. Alfredo Staffa and the Centre d’Expertise et de Services at Génome Québec for library preparation and RNA sequencing. We thank Julien Leconte and the Flow Cytometry and Cell Sorting at the Rosalind and Morris Goodman Cancer Research Centre for assistance with flow cytometry and single-cell analysis. We thank the McGill Integrated Core for Animal Modeling for housing of animal experiments described in this manuscript. We thank the Geneviève Morin and the McGill Centre for Cellular Perturbation for shRNA and overexpression constructs. We thank Valentina Muñoz Ramos, Camby Chhor, Nancy Laterreur, Monica Naujokas and the Rosalind and Morris Goodman Cancer Research Centre Research Support staff for administrative and logistical support. This work was supported by a Canada Foundation for Innovation (CFI) (33902), and a Canadian Institutes of Health Research foundation operating grants (242529) held by M.P. Y.S held a Charlotte and Leo Karassik Foundation Studentship, a Dr. Victor K.S. Lui Fellowship, a McGill Integrated Cancer Research Training Program Studentship, a Research Institute of the McGill University Health Centre Studentship, and a Canderel Studentship.

## AUTHOR CONTRIBUTIONS

Conception and design, Y.E.S., S.D., A.H.S.A., F.A., Z.Y., I.S., L.W. and M.P. Acquisition of data, Y.E.S., S.D., A.H.S.A. and A.M. Data analysis, Y.E.S., S.D., A.H.S.A., B.F. and L.W. Funding, M.P.

## COMPETING INTERESTS

The author have no competing interests to declare.

---

**Supplementary Table 1:**
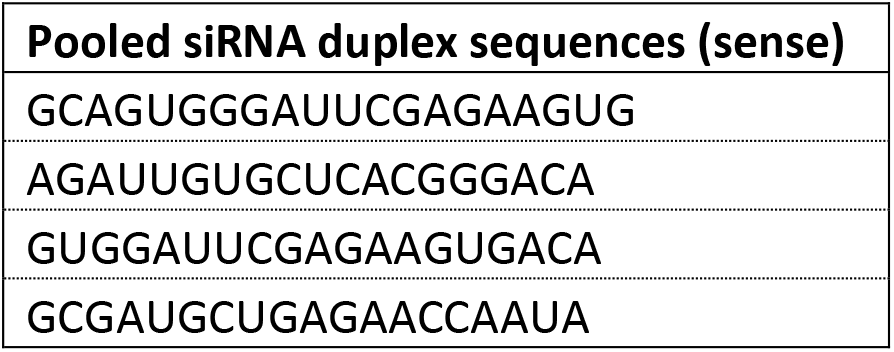
siRNA duplex sequences.

**Supplementary Table 2:**
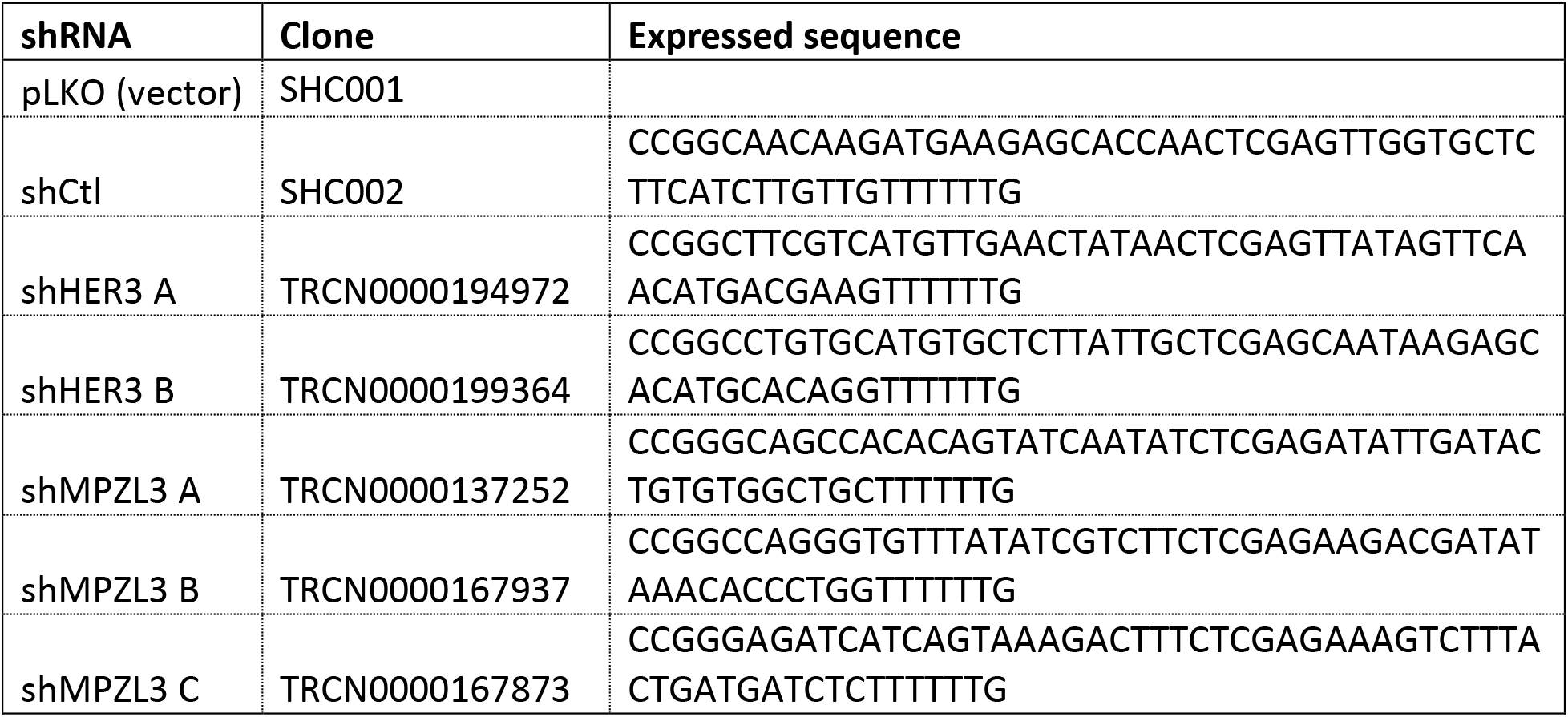
shRNA clone sequences.

**Supplementary Table 3:**
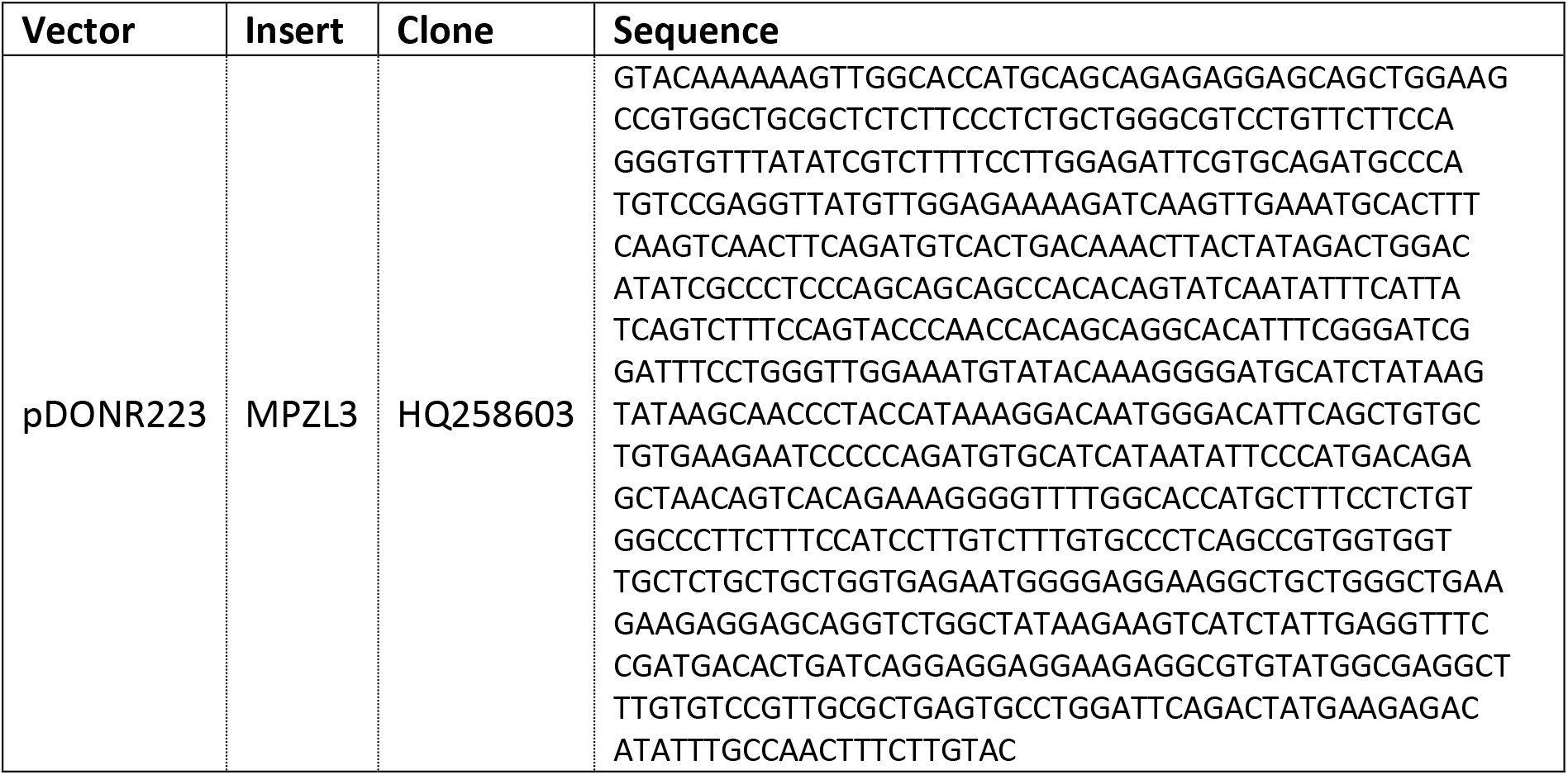
MPZL3 clone sequence.

**Supplementary Table 4:**
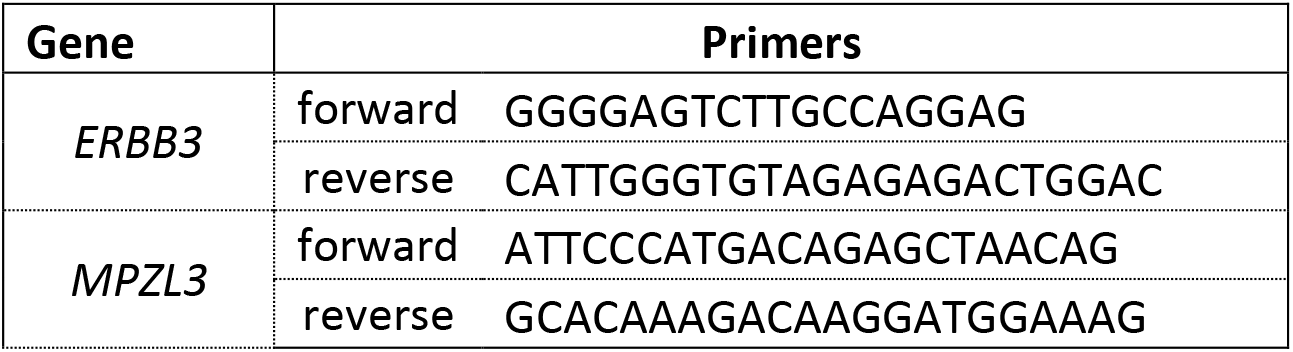
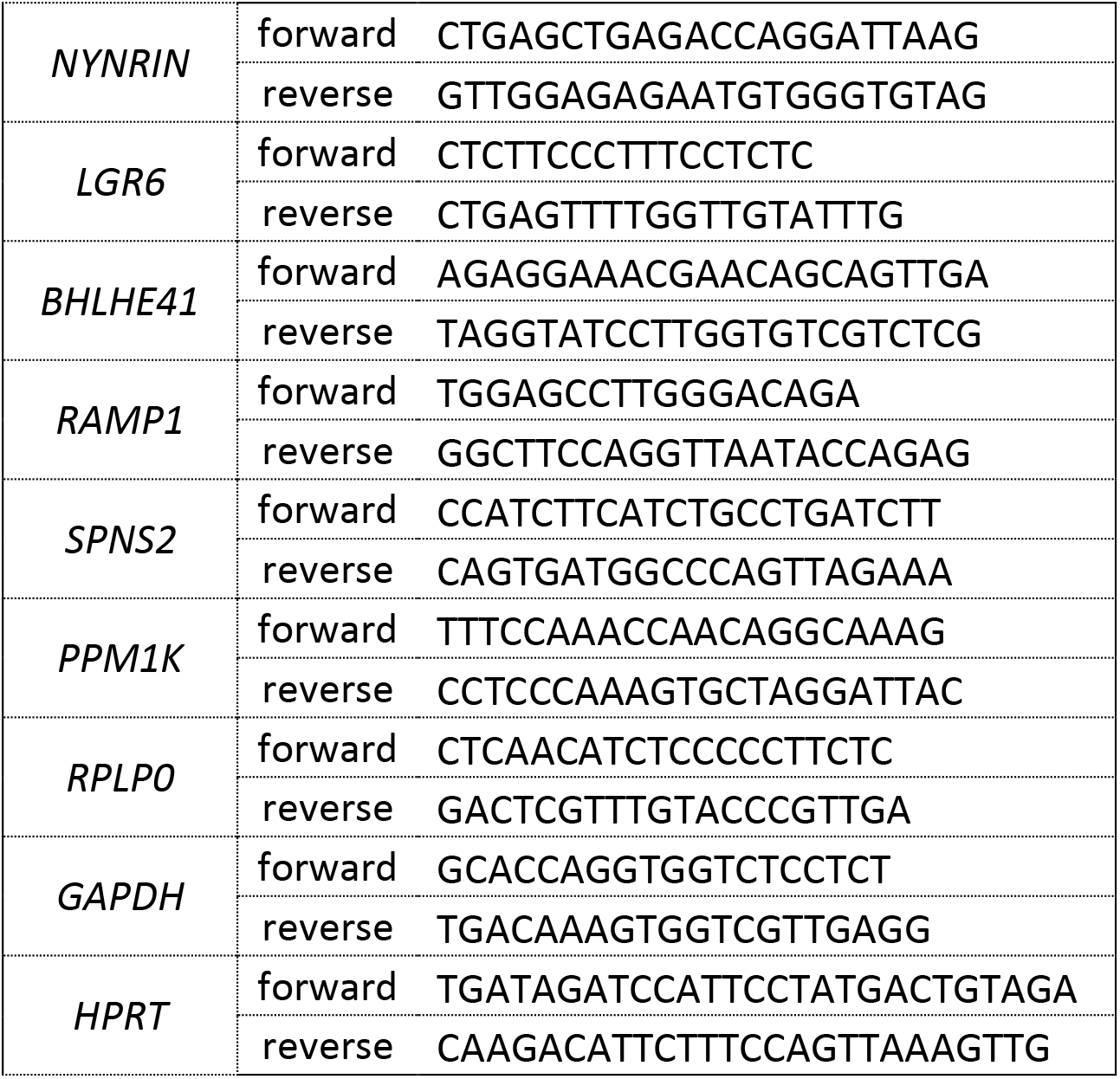
Sequences for RT-qPCR primers.

**Figure S1:**
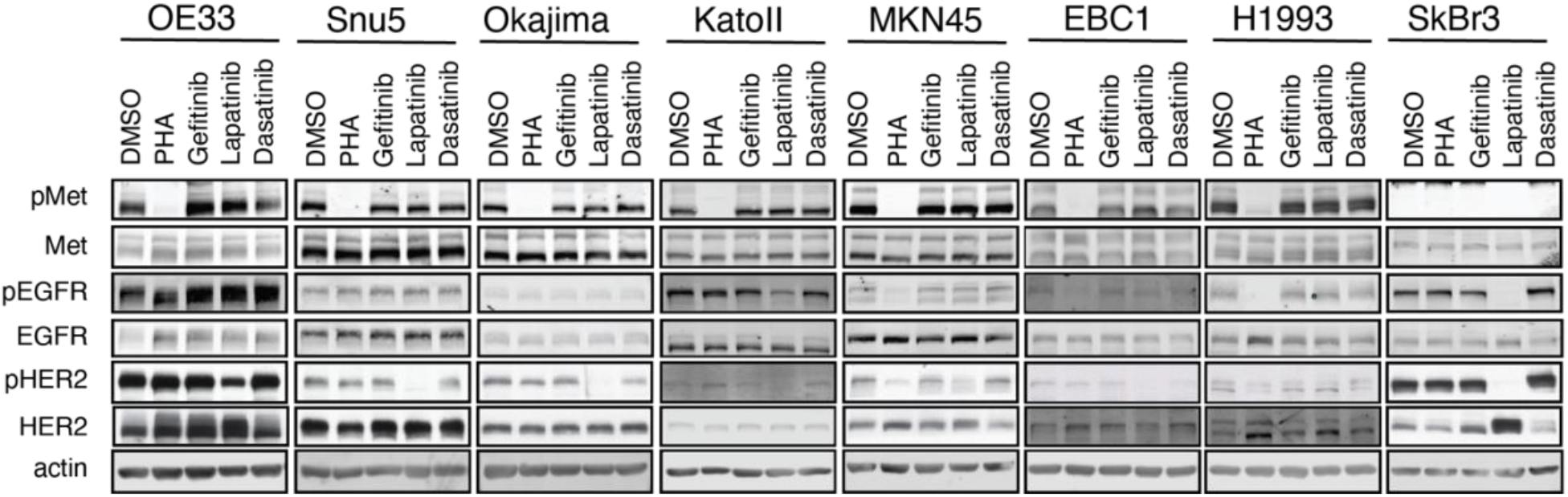
Controls for multiple kinase inhibitors used in *MET*-amplified cells (Figure 1C). Western blot analysis of the phosphorylation of Met (Tyr^1234/35^), EGFR (Tyr^1173^) and HER2 (Tyr^1221/22^) in the indicated cell lines treated with PHA (0.5 μM); the EGFR inhibitor, gefitinib (1 μM); the HER2/EGFR inhibitor lapatinib (1 μM); or the Src and Abl family kinase inhibitor, dasatinib (0.1 μM), for 1 hour (n=3).

**Figure S2:**
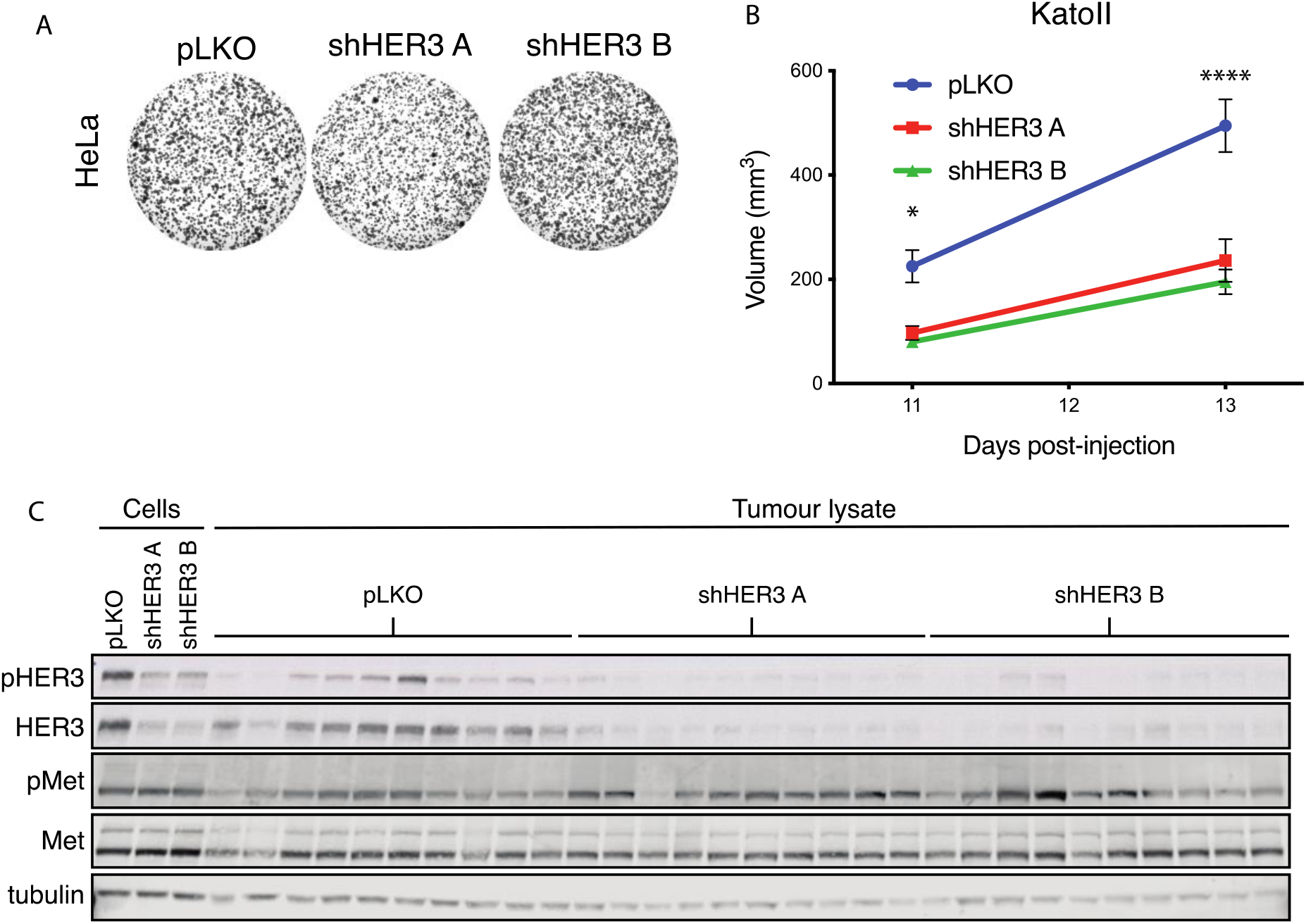
Growth of shRNA-treated cells in vitro and in vivo. (A) HeLa cells transduced with shRNA targeting HER3 form colonies at the same rate as controls (pLKO (n=3). (B) Tumour volumes of KatoII-derived xenografts shown in figure 2E at both points measured before collection (n=10). (C) Western blot analysis of total and phosphorylated Met and HER3 in KatoII tumours at endpoint.

**Figure S3:**
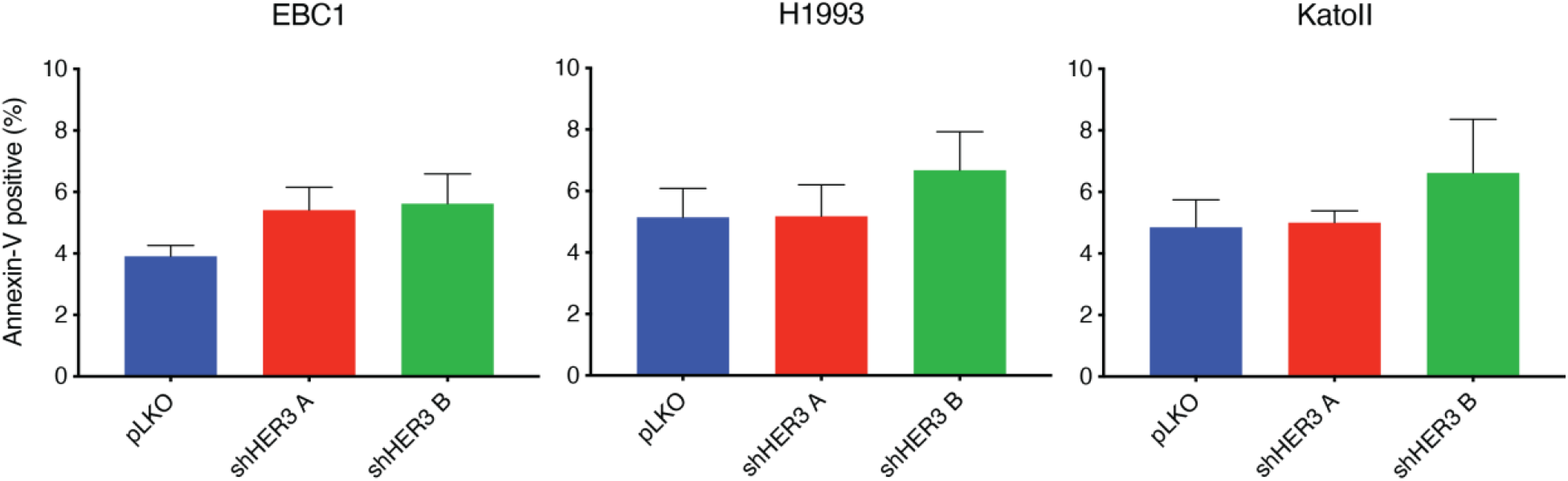
Apoptosis in shRNA-treated cells. Analysis of annexin-V positivity by flow cytometry in EBC1, H1993 and KatoII cells transduced with shRNA targeting HER3 or control (pLKO).

**Figure S4:**
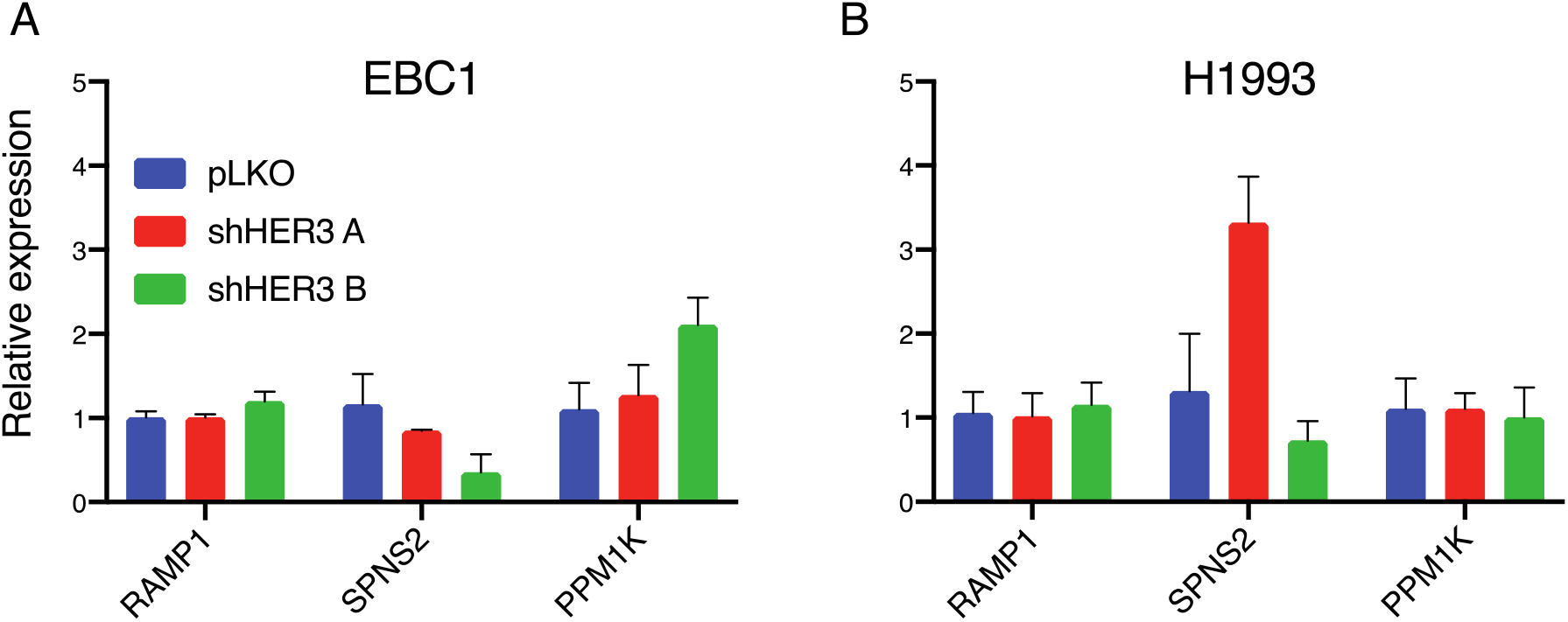
Gene expression analysis of upregulated transcripts from KatoII RNA sequencing in EBC1 and H1993 cells. Measurement of the relative amounts of RAMP1, SPNS2 and PPM1K genes in the indicated cell lines by RT-qPCR.

## Notes

### Competing Interest Statement

The authors have declared no competing interest.

